# Apoptosis promotes fertility in *C. elegans* by maintaining functional germline morphology

**DOI:** 10.1101/2025.10.22.683972

**Authors:** UN Saydee-Onwubiko, ME Werner, GC Trapp, AS Maddox

**Affiliations:** Department of Biology, University of North Carolina, Chapel Hill, NC 27599

**Keywords:** Apoptosis, Oogenesis, Germline architecture, Infertility, *C. elegans*

## Abstract

Programmed cell death (apoptosis) during oogenesis is conserved across metazoans and linked to regulation of oocyte number and quality. In oogenic germlines, the removal of developing oocytes by apoptosis ensures that oocytes do not contain DNA damage or multiple nuclei. Beyond this chromatin-quality control assurance role, it was unknown how apoptosis contributes to oocyte quality. We used the nematode *Caenorhabditis elegans* (*C. elegans*) to study the consequences of loss of apoptosis on oogenesis. Blocking apoptosis reduced fecundity in hermaphrodites at peak fertility and caused germline architectural defects such as abnormal rachis morphology and perturbed the arrangement and distribution of oogenic germline compartments. We posit that the defects associated with the loss of germline apoptosis arise due to lack of sufficient space for normal oogonia growth. In support of this idea, oocytes and embryos are abnormally small and exhibit low viability in animals unable to execute apoptosis. These findings suggest that in addition to preventing ploidy defects during oogenesis, apoptosis contributes to fertility by preserving homeostatic germline structure required for the fidelity of oogenesis.

**Summary Statement:** Blocking apoptosis during oogenesis disrupts germline structure over time, causing morphological defects and infertility, highlighting the role of apoptosis in maintaining syncytial architecture and reproductive function.

## Introduction

The US National Institutes of Health reported that one in four women of reproducing age experience infertility directly linked to oocyte quality defects, highlighting a need to understand the cellular mechanisms that ensure the fidelity of oocyte production or oogenesis (Nugent and Chandra, 2024; Pfennig et al., 2022). Oocyte quality (ploidy, size, and general physiology) is directly proportional to embryo fitness and viability (Conti and Franciosi, 2018; Hughes et al., 2018; Maddox et al., 2005). Cell size is of particular importance for oocyte function, since the egg must contain all biomass necessary for development until the embryo can access outside nutrients. The mechanisms that ensure oocyte quality is achieved and regulated are poorly understood (Frost et al., 2023; Luo et al., 2010).

Germ cell cysts in insects and vertebrates as well as oogenic germlines in nematodes, are syncytia (Hubbard and Greenstein, 2005; Spradling, 2024). Within these syncytial structures, germ “cells” are compartments in which germline nuclei reside, interconnected by actomyosin-rimmed cytoplasmic bridges (Edwards and Kiehart, 1996; Gerhold et al., 2022; Pepling and Spradling, 2001). In the germline, oogenic germ cells undergo mitotic divisions, become meiotic, and eventually experience one of two fates - elimination by apoptosis, or maturation into oocytes (Pepling, 2006; Pepling and Spradling, 2001). In all organisms examined to date, apoptosis during oogenesis occurs in germline compartments that are in the pachytene stage of meiosis I prophase (Du Pasquier et al., 2011; Kim et al., 2013; Niu and Spradling, 2022). Generally, apoptosis is essential for shaping tissues during development, and eliminating defective cells, which ensures homeostatic health and the avoidance of pathologies including cancer (Lindsten et al., 2000) During oogenesis in humans, mice, and *Drosophila*, apoptosis removes >90% of developing oocytes (Tilly, 2001). Mutations in apoptosis-related genes have been linked to maternal fertility defects in mammals (Ratts et al., 1995; Veis et al., 1993). In some species, apoptotic compartments serve a nurse cell function, providing cytoplasm to promote oocyte growth and development (McCall and Steller, 1998; Technau et al., 2003). In *Drosophila*, elimination of 15 of 16 oogonia by apoptosis is essential for oocyte enlargement and therefore viability (McCall and Steller, 1998). By contrast, in mammals, most growth of oocytes occurs after the breakdown of the germ cell cyst when apoptosis is prevalent (Anderson and Telfer, 2018; Niu and Spradling, 2022). In nematodes, cytoplasm from apoptotic compartments is thought to contribute a small amount of the massive volume of cytoplasm taken up by nascent oocytes prior to cellularization (Wolke et al., 2007); apoptosis has primarily been linked to ensuring elimination of compartments containing multiple nuclei or DNA damage (Raiders et al., 2018).

Apoptosis is initiated by the activation of proteases called caspases that degrade proteins to dismantle cellular structures (Kumar, 2007). The presence of multiple caspase pathways in most metazoans confounds deciphering the roles of apoptosis regulators in oogenesis (Khammari et al., 2011; McCall, 2004; Perez et al., 2007; Veis et al., 1993). In *C. elegans*, the effector caspase CED-3 is the sole caspase necessary and sufficient for germline apoptosis; loss of CED-3 is an elegantly simple way to experimentally tune apoptosis in this animal (Gumienny et al., 1999). The stereotypic and well-characterized development and function of the hermaphrodite germline also facilitate interrogating the roles of apoptosis in *C. elegans*. Hermaphrodites first generate ∼300 sperm, and transition into oogenesis in the fourth and final larval stage (L4) (Lints and Hall, 2009; Pazdernik and Schedl, 2013). At 24 hours post L4, animals are young adults, and their germlines make oocytes that are fertilized via sperm stored in the spermatheca (Pazdernik and Schedl, 2013). The oogenic germline consists of two U-shaped tubes that each connect, via a spermatheca, to a central uterus (Kimura and Motegi, 2025). In a stem cell niche in the distal tip of the germline, germline stem cells undergo mitotic proliferation (see Fig. 1C). When they are displaced out of the niche, germline compartments transition into meiosis. Approximately 50% of compartments in pachytene of meiosis I prophase undergo apoptosis (Gumienny et al., 1999), while the rest exit pachytene in a Ras/MAP kinase-dependent manner, progressing through diplotene and diakinesis until they receive signals to complete maturation and be ovulated (Church et al., 1995; Huelgas-Morales and Greenstein, 2018; Lee et al., 2007b). Once ovulated from the uterus and fertilized, oocytes complete meiosis and begin embryogenesis (Hubbard and Greenstein, 2005). *C. elegans* hermaphrodites’ period of fertility depends on the presence of sperm, which is finite in the absence of mating; the exhaustion of the sperm supply marks post-fertility (McCarter et al., 1999; Scharf et al., 2021).

**Figure 1.**
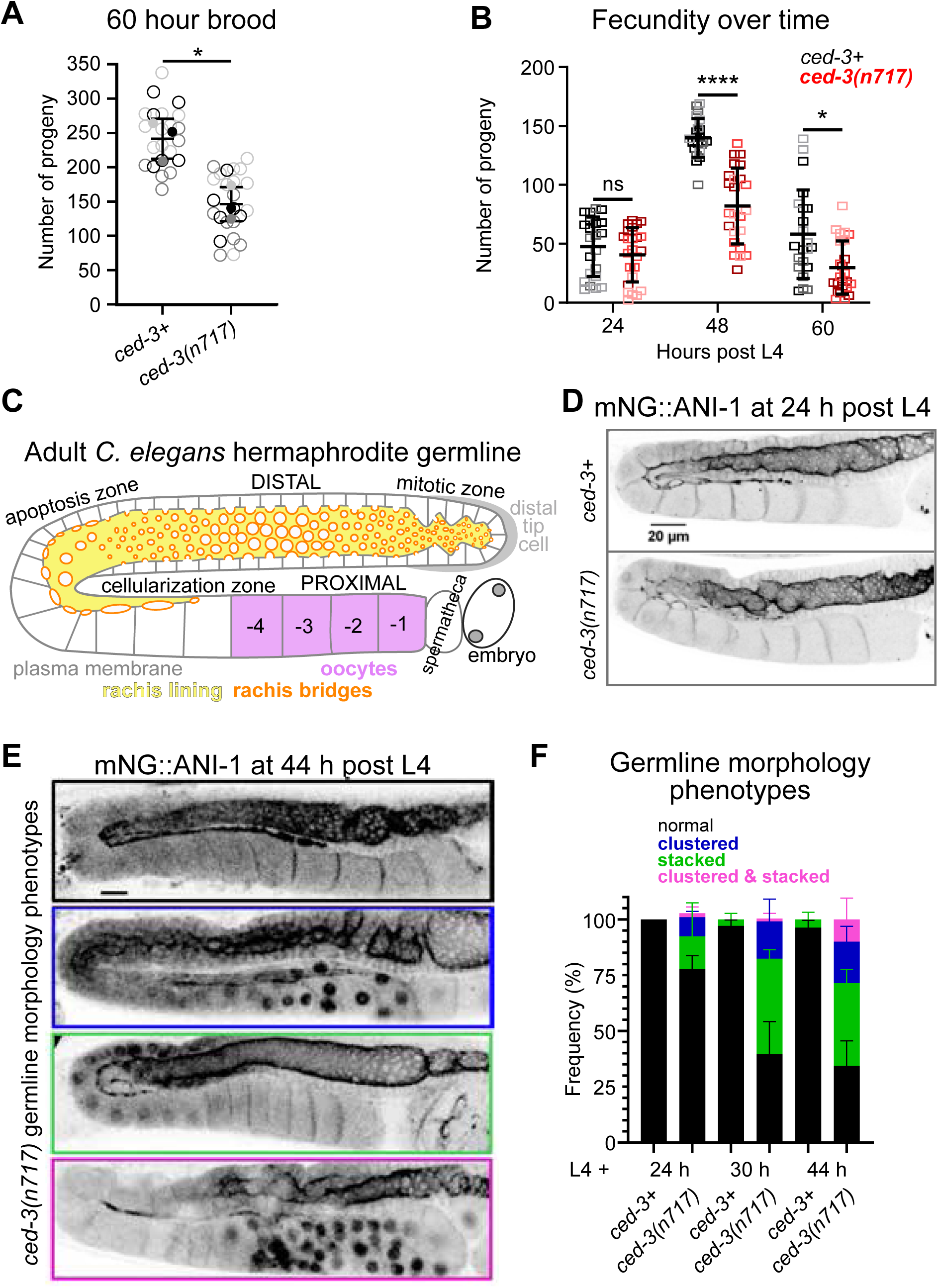
*ced-3(lf)* hermaphrodites exhibit brood size and germline architecture defects that intensify over the reproductive lifespan. **A.** Quantification of total brood size in the indicated genotypes. Data points represent individual worms from 3 independent experiments (various gray levels) n≥10 hermaphrodites per condition analyzed). Statistical significance determined by Welch’s Two-tailed t-test, * p-value = 0.01. **B.** Quantification of average brood size in time windows 0-24, 24-44, and 44-60 hours post larval stage 4 (L4). Data points represent individual worms from 3 independent experiments (n = 24 animals per genotype). Statistical significance determined by ANOVA followed by Šídák’s post hoc analysis, n.s (not significant, p=0.708), * p-value = 0.01, **** p<0.0001.**C.** Schematic of an adult *C*. *elegans* hermaphrodite germline highlighting key morphological features and zones. **D.** Images of control and *ced-3*(*lf*) germlines at 24 hours post L4 from animals expressing mNG::ANI-1 from its endogenous locus (MDX29 and MDX129). **E.** Representative images of germlines in animals expressing mNG::ANI-1 of germline morphologies scored in F: black (normal), blue (clustered germ compartments), green (stacked oocytes), and magenta (clustered germ compartments + stacked oocytes). **F.** Quantified frequency of germline morphologies of control and *ced-3*(*lf*) animals at different reproductive time points (L4+24h, L4+30h, and L4+44h, 3 independent experiments, n ≥ 80 animals per experimental condition. Error bars: mean ± s.d., Scale bars: 20 µm in all panels.

The *C. elegans* caspase CED-3 is activated through physical interaction with the pro-apoptotic Apaf-1 homolog CED-4 (Hengartner et al., 1992; Wu et al., 1997). Mutations in *ced-3* or *ced-4* that alter their physical interaction phenocopy each other (Gumienny et al., 1999; Hengartner et al., 1992). How CED-3-driven apoptosis affects oocyte quality is not understood. On one hand, aged, post-fertility *ced-3*(*lf*) mutants have been reported to exhibit germline hyperplasia (Gumienny et al., 1999) and reduced oocyte quality (Andux and Ellis, 2008). However, other work reports that *ced-3*(*lf*) animals are viable, fertile, and have normal germline architecture when actively producing oocytes that are fertilized (peak fertility) (Das et al., 2020). To address this conundrum and understand the cellular implications of apoptosis in oogenesis, we tested whether blocking apoptosis perturbs germline morphology and reproductive success, across several developmental timepoints. We discovered that blocking apoptosis reduced maternal fertility much earlier than previously reported (Andux and Ellis, 2008; Gumienny et al., 1999), within the timeframe of normal fertility. Apoptosis-defective germlines exhibited morphological defects including abnormal rachis architecture, reduced cross-sectional surface area of germline compartments, and displacement of compartments from their normal positioning within the germline. *ced-3(lf)* animals yielded abnormally small embryos; these small embryos had decreased embryonic viability. These findings provide insights into the functional relevance of apoptosis and germline morphology for fertility.

## Results

### *ced-3(lf)* hermaphrodites exhibit brood size and germline architecture defects that intensify over the reproductive lifespan

Apoptosis has been linked to maintenance of germline architecture and disassembly in aged hermaphrodites exhausted of sperm (de la Guardia et al., 2016; Gumienny et al., 1999). Additionally, loss of apoptosis was also linked to increases in age dependent embryonic lethality in mated virgin females (Andux and Ellis, 2008). Finally, slight decreases in embryonic viability and total brood size have also been observed in apoptosis defective animals (Ellis and Horvitz, 1986; Hengartner et al., 1992). However, no cell biological characterization of the origin of these defects in fertile hermaphrodites has been carried out to date. We used the apoptosis-defective *ced-3* loss of function mutant *ced-3*(*n717*) (“*ced-3(lf)”*) (Ellis and Horvitz, 1986) to recapitulate previously reported defects in hermaphrodite fertility (Ellis and Horvitz, 1986; Hengartner et al., 1992). The total number of embryos laid (brood size) within the fertile period (the 60 hours following the fourth larval stage (L4)) was significantly lower for *ced-3*(*lf*) hermaphrodites than wild type control animals (Fig. 1A). Given the age-dependent effect of loss of apoptosis on embryonic viability in mated virgin females, we tested whether defects in fertility showed an age dependent progression in severity in fertile hermaphrodites as well. We measured the average brood count in three distinct time windows: 0-24, 24-44, and 44-60 hours post L4. At 24 hours, control and *ced-3(lf*) brood sizes were comparable; however, the mutants failed to maintain a normal brood size from that point onwards (Fig. 1B). The reduction in brood size, and therefore germline function, in *ced-3(lf)* animals suggested that apoptosis is required for normal oogenesis.

Various aspects of germline morphology are correlated with successful oogenesis (Green et al., 2011). Young adult *ced-3(lf)* mutant germlines are reported to be morphologically indistinguishable from wild type, while post-fertility aged germlines (72 hours post L4) show defects in germline architecture (Das et al., 2020; Gumienny et al., 1999). To test whether *ced-3*(*lf*) fertility defects in older fertile hermaphrodites were due to changes in germline morphology, we examined germlines of animals expressing fluorescently labelled anillin-1 (ANI-1), a cortical scaffolding protein (Maddox et al., 2005; Rehain-Bell et al., 2017). ANI-1 localizes to the germline cortex, rachis bridges, oocyte boundaries, and rachis lining (Amini et al., 2014; Rehain-Bell et al., 2017) (Fig. 1C, D). In accordance with others’ and our observation that *ced-3(lf)* mutants generated a normal number of progeny in the first 24 hours of adulthood, germline architecture was largely normal 24 hours after L4 (Das et al., 2020) (Fig. 1D). 20 hours later (44 hours into adulthood), *ced-3*(*lf*) mutant animals laid significantly fewer progeny (Fig. 1B), and germlines were morphologically abnormal (Fig. 1E and F). While in control germlines, developing oocytes were arranged in a single row throughout the spermatheca-proximal germline (Fig. S1A), in *ced-3*(*lf*) germlines, oocytes in the proximal germline were often abnormally small and failed to form a single row (“clustered”), abnormally numerous and occupying a single row of short cylinders (“stacked”), or both (Fig. 1E and F). Since accumulation of “stacked” oocytes can occur when hermaphrodites are depleted of sperm (Tolkin and Hubbard, 2021), we assessed the time course of the appearance of the various phenotypes. *ced-3(lf)* germlines first exhibited stacked, then clustered, and a combination of both, as animals progressed from 24 to 44 hours post L4 (Fig. 1F). Approximately 5% of control animals exhibited stacked oocytes in the proximal germline at 44 hours post L4, but no other defects were observed (Fig. 1F). The percentage of *ced-3(lf)* mutant germlines that appeared normal decreased over time to ∼33% at 44 hours post L4 (Fig. 1F). Thus, morphological anomalies develop before exhaustion of the sperm supply and intensify as animals age. These observations collectively highlight the importance of germline apoptosis in ensuring normal germline morphology.

### *ced-3(lf)* germlines have abnormal rachis and germline compartment morphology

To better understand how loss of apoptosis contributes to defects in germline architecture, we first assessed the effects of *ced-3(lf)* on the number and organization of germline compartments. We introduced an mCherry-tagged membrane-binding probe into *ced-3(lf)* to label germline membranes and documented both germline morphology and germline compartment number at 44 hours post L4 when significant germline architecture defects were observed (Fig. 2A). Germline compartments of control animals formed a single row around the germline bend and throughout the proximal germline. By contrast, these germline domains in *ced-3(lf)* animals were occupied by smaller, often irregularly shaped compartments (Fig. 2A). We then quantified the number of germline compartments in each 10 µm bin throughout the germline of both control and *ced-3(lf)* animals 44 hours post L4. The number of germline compartments per unit length was consistently higher in *ced-3*(*lf*) mutants than in controls starting at 100 µm distal to the bend in the germline which in wild type worms corresponds to mid to late pachytene, the region where most germ cell apoptosis occurs (Fig. 2B, B’). These observations suggested that the absence of apoptosis leads to an increase in the overall number of germ cells occupying the germline leading to defects in overall germline architecture.

**Figure 2.**
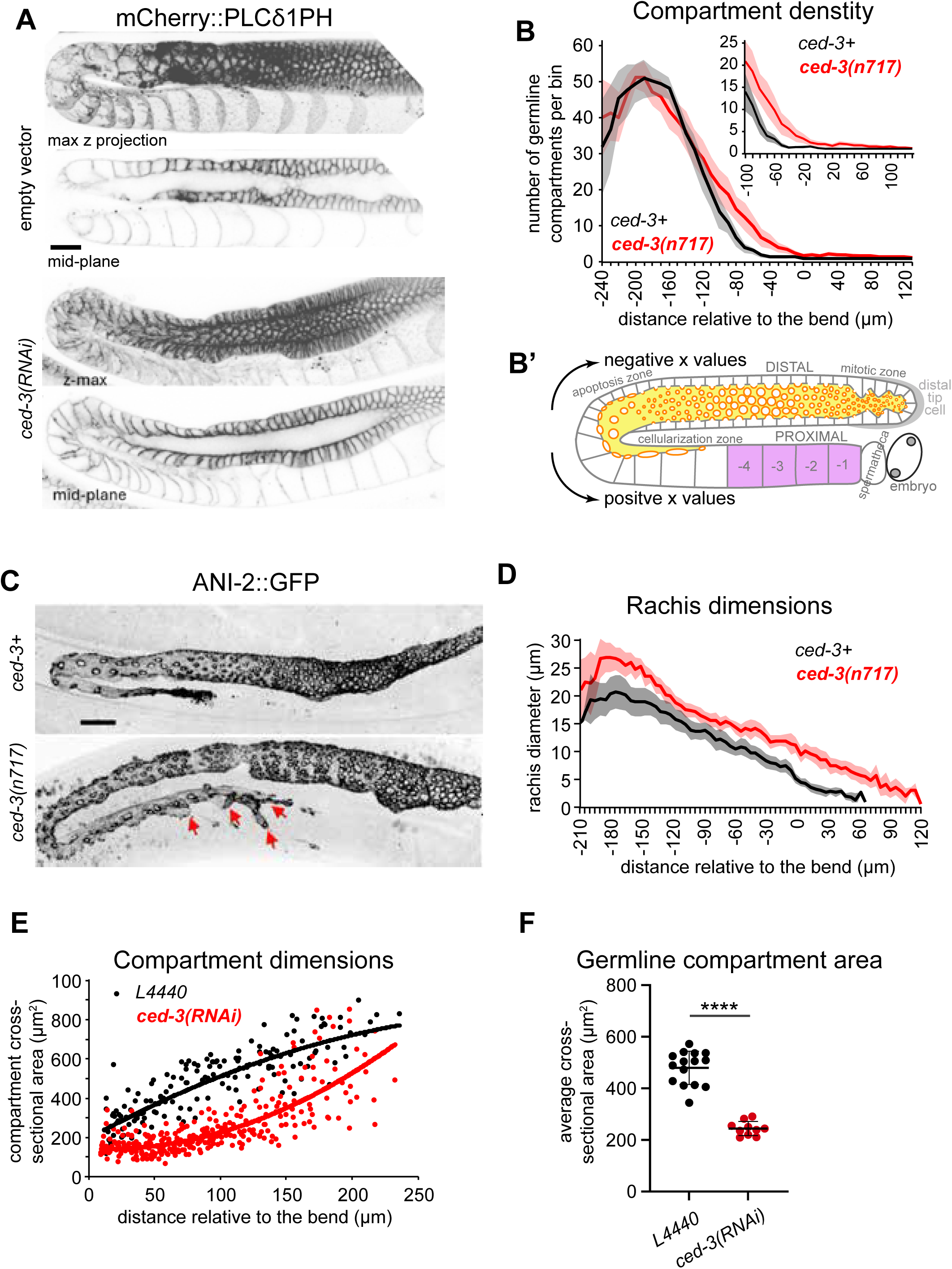
*ced-3(lf)* germlines have abnormal rachis and germline compartment morphology. **A.** Representative images of membrane (mCherry::PH, MDX63) localization used for quantifications in B and E. Animals were fed with RNAi bacteria for 44hours; L4440: empty vector (top panels), and *ced-3(RNAi)* (lower panels). **B.** Average number of compartments within 10 µm bins of the germline, at positions relative to the bend in control (n=14) and *ced-3(n717)* (n=20) mutant worm. Shaded areas represent 95% confidence intervals. B’ Schematic depicting position of positive and negative germline distances (x-values) from bend, represented in B, D and E. C. Representative images (max-projections) of the rachis (ANI-2::GFP) in the indicated genotypes at 44 hours post L4, red arrows highlight abnormal bifurcations/protrusions. **D.** Quantification of rachis diameter throughout the germline (from distal tip to cellularization) in the indicated genotypes, n=14 animals germlines per genotype. Shaded areas represent 95% confidence intervals. **E.** Cross-sectional area of individual compartments in the proximal germline for indicated genotypes. **F.** Quantification of average cross-sectional area of germline compartments in control and CED-3 depleted animals. ****: p < 0.0001 Student’s two-tailed t-test. For E and F, animals were imaged at 45 hours post L4 on RNAi bacteria plates, n = 15 control (L4440), and 10 *ced-3(RNAi)* germlines. All scale bars: 20 µm.

Germ cell compartments within the syncytial germline are connected to the common cytoplasm (the rachis) that acts as a net acceptor or donor of cytoplasm depending on the position within the germline (Maddox et al., 2005; Nadarajan et al., 2009; Wolke et al., 2007). In wild type animals, near the germline bend, the rachis becomes a net donor of cytoplasm and shrinks as surrounding germline compartments enlarge, suggesting that rachis morphology plays an important role in overall germline architecture and oogenesis (Chartier et al., 2021) (Fig. 2B’). To test whether changes in germline compartment number in *ced-3(lf)* germlines correlated with abnormal rachis morphology, we measured rachis diameter throughout the germline using ANI-2::GFP, which specifically marks the rachis lining, in control and *ced-3*(*lf*) animals 44 hours post L4. In *ced-3*(*lf*) germlines as in controls, the rachis steadily narrowed from pachytene to cellularization but was wider along its entire length in *ced-3*(*lf*) animals (Fig. 2C, D). Further, 90% (45/50) of *ced-3* mutant germlines displayed a bifurcated rachis at 44 hours post L4, compared to 0% (0/52) in control animals (red arrows, Fig. 2C).

To determine if the changes in germline architecture (increased density of germline compartments and increased rachis diameter in *ced-3(lf)* mutants), translated into changes in germ cell and oocyte size we next measured the cross-sectional area of individual compartments proximal to the bend at 44 hours post L4 in both control and *ced-3(lf)* worms. In *ced-3*(*lf*) mutant animals, germline compartments past the germline bend were smaller compared to controls (Fig. 2E). On average, the cross-sectional area of control compartments was two-fold that of the mutants, underestimating the difference in compartment volume (Fig. 2F). Notably, the compartments furthest from the bend (the cellularized oocytes) were also often markedly smaller in *ced-3(lf)*, although the difference in size was not as pronounced for germ cell compartments more proximal to the bend; some oocytes’ size was comparable to controls (Fig. 2E). Taken together, these results suggested that loss of apoptosis caused an increase in overall germline compartment number, which leads to changes in germline architecture characterized by a widening of the rachis and a decrease in germline compartment and oocyte size.

### *ced-3(lf)* mutants compensate for excess germ cells by delaying cell cycle progression and cellularization

Given the significant changes to germline architecture in the absence of germline apoptosis, it was notable that overall fertility and embryonic viability are only slightly reduced in *ced-3(lf)* worms (Ellis and Horvitz, 1986; Hengartner et al., 1992). This raised the question of how animals compensate for the increase in germline compartment number and corresponding decrease in their size. The most straightforward possibility for accommodating the increase in germline compartment number would be to extend the length of the germline. We therefore measured the total length of the germline from distal tip to the spermatheca in control and *ced-3(lf)* animals 44 hours post L4. Germline overall length was slightly longer in *ced-3(lf)* suggesting that mutants have a limited ability to increase germline size in response to an increase of germline compartment number (Fig. 3A).

**Figure 3.**
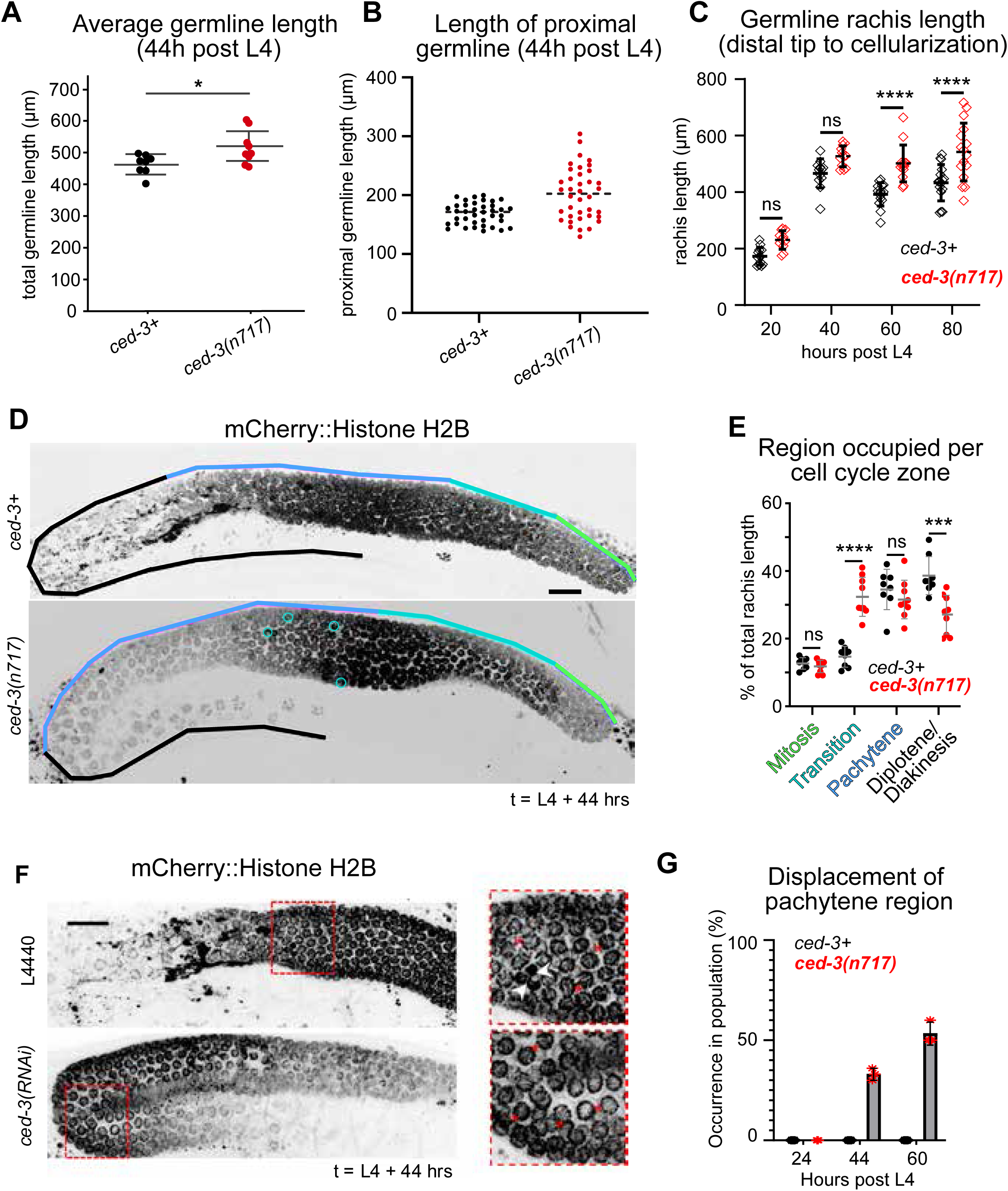
*ced-3(lf)* mutants compensate for excess germ cells by delaying cell cycle progression and cellularization. **A.** Measurements of total germline length from distal tip to the spermatheca in control (MDX155, n=8) and ced-3 mutant animals (MDX187) at 44h past L4. Statistical significance was determined using a 2 tailed student T-Test, p<0.05. **B.** Quantification of the length of the proximal germline measured from the bend to cellularization in the indicated genotypes, at 44 hours post L4; n=38 germlines per condition. Error bars: mean ± s.d; statistical significance determined by Welch’s Two-tailed t-test, **** p-value ≤ 0.0001. **C.** Quantification of germline rachis length in synchronized populations in the indicated genotypes, [n ≥ 13 germlines per condition, and time post L4 indicated. Error bars: mean ± s.d. Statistical significance determined by 2-way ANOVA with Šídák’s multiple comparisons post-hoc test, ****p ≤ 0.0001, n.s.: not significant, p=0.092, and 0.062]. **D.** Images of germline nuclei (H2B::mCherry) organization in the mentioned genotypes at 44h post L4, with colored lines indicating cell cycle stages of germline nuclei based on chromatin appearance, cyan circles highlight intermingled transition state nuclei within pachytene. Scale bar: 20 µm. **E.** Quantification of the percentage of the rachis occupied by mitotic, transition, pachytene and diplotene/ diakinesis nuclei in *ced-3*+ and ced*-3*(*lf*) germlines, at 44 hours post L4, n = 7 - 8 germlines per genotype analyzed. [Statistical significance determined by Two-way ANOVA with Tukey’s multiple comparisons test, **** p-value ≤ 0.0001, *** p-value ≤ 0.001, n.s.: not statistically significantly different]. **F.** Max-projection images of germ compartment nuclei (H2B::mCherry) in the experimental conditions indicated. Pachytene nuclei (red dashed box). Within the enlarged images of nuclei (insets), apoptotic nuclei in an L4440 control germline (white arrows), pachytene nuclei displaced in *ced-3*(*RNAi)* conditions (Red asterisks) **G.** Quantification of displacement of pachytene nuclei in the indicated genotypes. Data represent three blindly scored independent experiments (n = 10 germlines per genotype). Error bars: mean ± s.d. in all panels unless specified otherwise. All scale bars: 20 µm.

Another way to promote proper oogenesis could be to delay cellularization of oocytes to increase the time for individual compartments to accept cytoplasm from the rachis and reach the appropriate size. To test this possibility, we measured the length of the proximal rachis (from the distance from the bend to the end of the rachis) in control and *ced-3(lf)* animals 44 hours post L4. In *ced-3(lf)* mutants or following CED-3 depletion by RNAi, the proximal rachis was longer than in control animals and often appeared bifurcated at its end (red arrows, Fig. 2C, D, 3B and S1B). This effect arose only after 24 hours following the L4 stage (Fig. 3C). This suggests that in the absence of apoptosis, delaying cellularization may be one way to assure viable oocyte formation.

Oogenesis depends on the progression of germline compartments from mitosis into, and through the stages of prophase of meiosis I (Kim et al., 2013). The number of compartments in each stage (and thus the length of the germline they occupy) reflects the duration of each stage. Another way to compensate for an excess number of germline compartments accumulated due to lack of removal by apoptosis would be to slow cell cycle progression within the germline. To explore this possibility, we assessed DNA morphology using an mCherry-tagged histone variant (H2B::mCherry) (Fig. 3D). Chromatin morphology allows for the reliable identification of meiotic stages, as defined previously (Crittenden et al., 1994; Hubbard, 2007; Jaramillo-Lambert et al., 2007; Lints and Hall, 2009). We classified the nuclear morphology of compartments in control and *ced-3(lf)* mutant animals 44 hours after L4. In control animals, compartments immediately preceding the bend were in pachytene, and compartments around the bend and in the proximal germline were in first diplotene and then diakinesis; groups of compartments in each stage were distinct and did not intermingle (Fig. 3D). In *ced-3(lf)* mutants, the transition zone was extended, and some leptotene/zygotene nuclei intermingled with pachytene nuclei (Fig. 3D cyan circles). In some cases, the pachytene zone extended around the bend and into the proximal arm of the germline (Fig. 3F). Measurements of the proportion of the germline rachis occupied by mitotic, transition, pachytene, and diplotene/diakinesis nuclei revealed that the fractions occupied by mitotic and pachytene compartments were similar in controls and *ced-3(lf)* mutants (Fig. 3E). By contrast, the transition zone of *ced-3(lf)* mutants occupied a larger fraction of the germline than those of controls (Fig. 3E), and this was only observed 44 and 60 hours post L4 (Fig. 3G). This suggests that the reduced elimination of compartments in pachytene is compensated for by a delayed exit from the transition zone in *ced-3*(*lf*) animals.

Taken together, our findings suggest that loss of apoptosis leads to delays in cellularization and cell cycle progression that partially compensate for the defects in germline architecture and decreases in germ cell volume that accompany increased germ cell number. Thus, germline apoptosis may contribute to oogenesis by controlling the rate of compartments’ transit through meiosis and enlargement.

### Apoptosis promotes embryonic viability by ensuring oocyte size

It has been reported that loss of apoptosis leads to a decrease in embryonic viability (Ellis and Horvitz, 1986; Hengartner et al., 1992). This could be due to the failure to remove germline compartments with DNA damage and multinucleation, or due to defects in embryo size as a consequence of defects in germline architecture, or a combination of both. To explore these possibilities, we assessed the viability of embryos produced by control and *ced-3*(*lf*) animals over time. Compared to controls, the viability of *ced-3(lf)* mutant embryos was 20% lower, even in early fertility (0-24 hours post L4) when only mild germline architecture defects were observed. This likely represents the baseline of embryonic lethality due to the fertilization of oocytes with DNA damage or multinucleation (Fig. 4A; (Raiders et al., 2018)). In the next 24 hour time window (24-44 hours post L4), at the end of which germline architecture defects are readily observed, embryonic lethality is the same as in earlier stages. By contrast, at 44-60 hours post L4, embryonic lethality was significantly higher, reaching nearly 50% (Fig. 4A). These data suggested that at least some of the observed defects in embryonic viability are due to defects in germline architecture.

**Figure 4.**
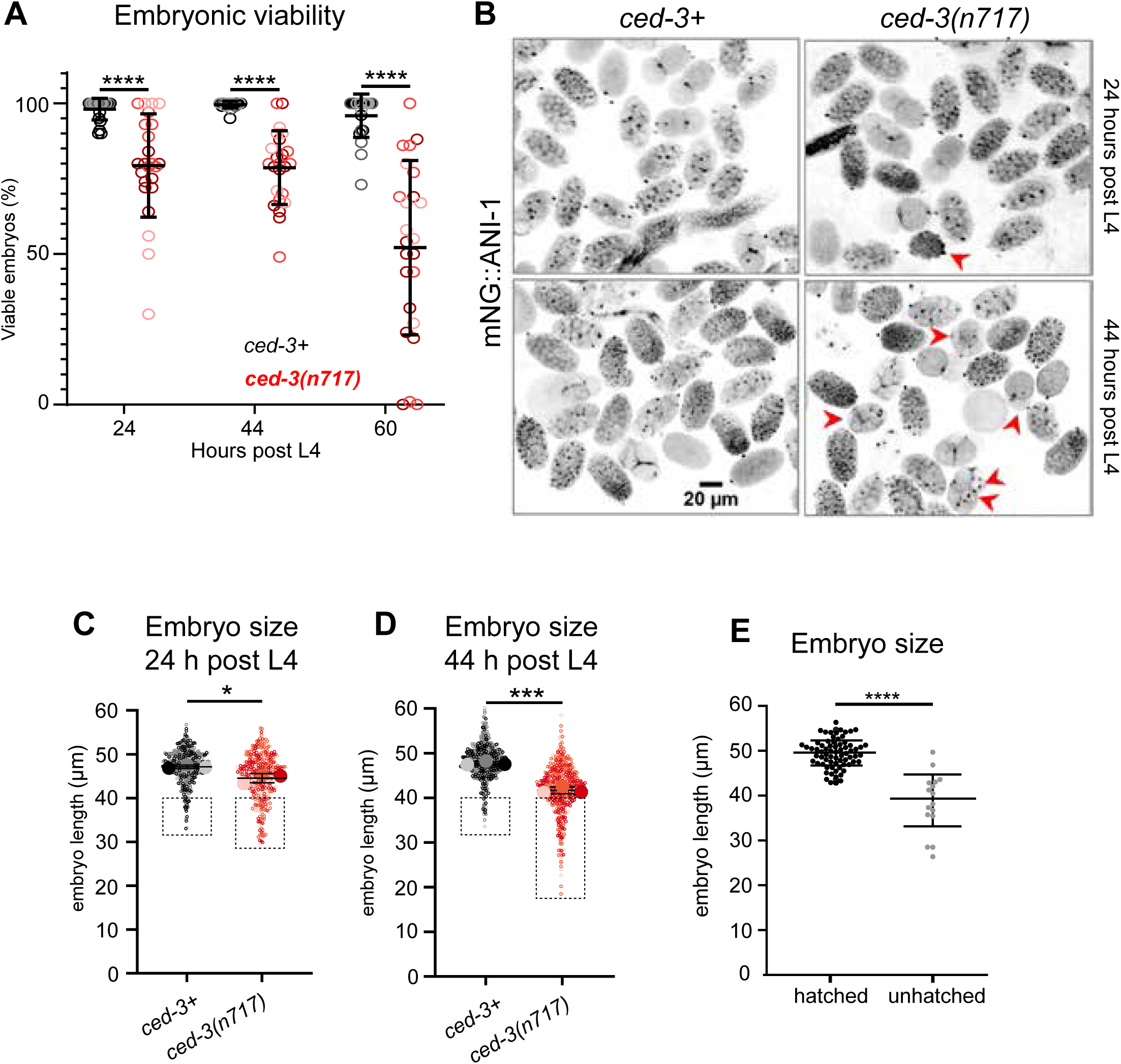
Apoptosis promotes embryonic viability by ensuring oocyte size. **A.** Embryonic viability of progeny of animals at 0-24, at 24-48 and 48-60 hours post L4, of control and *ced-3(n717)* animals. Data points represent individual worms from 3 independent experiments (n = 24 animals per genotype). Statistical significance determined by ANOVA followed by Šídák’s post hoc analysis, ****p < 0.0001, Error bars: mean ± s.d,). **B.** Representative images of embryos extruded from animals at 24 and 44 hours post L4 in the indicated genotypes. Red arrows point to abnormally small embryos. Scale Bar: 20 µm. **C & D.** Quantification of embryo length in the listed genotypes for animals at 24 hours post L4. Grey percentages: percent embryos smaller than 40 µm long. Three independent experiments, each with n ≥14 animals per genotype. Statistical significance determined by Welch’s Two-tailed t-test, * p= 0.0361, ***p= 0.0009. Error bars: mean ± s.d, **E.** Quantification of embryo length and viability from overnight timelapse imaging of embryos pooled from *ced-3*(*lf*) animals at 44 h post L4, n = 15 animals. Statistical significance was determined using a 2 tailed student T-Test, p<0,0001.

Given that smaller embryos are less likely to be viable (Hubatsch et al., 2019; Maddox et al., 2005) and that one of the major defects we observe in *ced-3(lf)* animals is a significant reduction in germline compartment and oocyte size (Fig. 2E,F), we asked if the increase in embryonic lethality observed 44 hours and longer after L4 correlated with an increase in the population of small embryos. We measured embryo size by extruding embryos from gravid hermaphrodites at 24 and 44 hours post L4, to assess embryo populations in the 24-44 and 44-60 hours post L4 time windows, respectively. Compared to wild type embryos that appeared normal in size, many *ced-3(lf)* embryos were abnormally small at 44 hours post L4 (Fig. 4B). Specifically, ∼35% of *ced-3(lf)* embryos were at a length of 40 µm or less compared to ∼5% of control embryos. Small embryos were also produced, rarely, by *ced-3(lf)* animals at 24 hours post L4 (Fig. 4C, D), possibly reflecting how some *ced-3*(*lf*) animals displayed morphological defects of stacking and clustering even at 24 hours post L4 (Fig. 1F).

To confirm that small *ced-3(lf)* embryos are indeed those that fail to develop into viable worms, we dissected *ced-3*(*lf*) animals at 44 hours post L4 to observe their embryos by time-lapse imaging for 24 hours at 20° C. Whereas wild type *C. elegans* complete embryogenesis and hatch within 12 hours of fertilization at 20° C (Hall et al., 2017), 19% of *ced-3*(*lf*) embryos failed to hatch (Fig. 4E) even after 24 hours. We measured the size (anterior-posterior axis length) of each embryo and scored whether hatching occurred. Embryos that hatched were on average 49 µm in length while the average length of unhatched embryos was 39.5 µm. All *ced-3*(*lf*) embryos over 40 µm in length hatched; all those below 40 µm in length were unable to complete embryogenesis (Fig. 4E). Some normal sized embryos died as well, potentially reflecting the subset of embryos dying as a consequence of DNA damage or multinucleation (Fig. 4E). This suggests that increased embryonic lethality in *ced-3(lf)* 44 hours post L4 animals is indeed caused by reduced embryo size, which itself arises from defects in germline architecture. These findings indicate that in addition to ensuring embryonic viability via DNA damage surveillance in the germline, apoptosis also contributes to embryonic fitness by maintaining germline morphology, to accomplish the production of viable progeny.

## Discussion

We investigated how apoptosis promotes fertility and oocyte quality during oogenesis in *C. elegans* using loss-of-function perturbations of the germline apoptosis caspase CED-3. We find that worms lacking apoptosis exhibit fertility defects that become more pronounced with age. The observed reduction in brood size correlates with an age dependent increase in defects in germline architecture. These defects included an increase in germline compartment number, an increase in rachis cross-sectional area, and a decrease in germ cell size proximal to the bend. These defects may be partially counterbalanced by the observed increase in the duration of the transition zone stage and extension of the proximal rachis to delay cellularization. The appearance of architectural defects coincided with an age-dependent increase in embryonic lethality that likely reflects fertilization of smaller oocytes that fail to develop.

Our time course analysis of brood size, embryonic viability, and germline architecture allowed us to reconcile seemingly contradictory results from previous studies that reported no significant germline defects in young adult animals (Das et al., 2020; Gumienny et al., 1999) and reduced brood size and embryonic viability, as well as significant germline architecture defects, in hermaphrodites exhausted of sperm (Ellis and Horvitz, 1986; Gumienny et al., 1999; Hengartner et al., 1992). We propose that in the absence of apoptosis, *C. elegans* animals partially compensate for the increase in total germline compartments that are no longer removed during pachytene by delaying exit of germ cells from the transition zone and extending the proximal rachis. By extending the area occupied by transition zone compartments, the onset of diplotene and diakinesis, when compartments significantly increase in size (Rehain-Bell et al., 2017) is delayed, by extension delaying the enlargement and cellularization of oocytes. These compensatory mechanisms may be sufficient to ensure normal oocyte morphology and number in young adults (Das et al., 2020; Gumienny et al., 1999) the effect on overall brood size and embryonic viability. As the animals age, however, the germline loses the capacity to compensate for the excess compartments, and defects in germline architecture, fecundity, and embryonic viability result (as reported by (Ellis and Horvitz, 1986; Hengartner et al., 1992). An open question in this context remains: how is this spatial adaptation of meiotic progression coordinated with cell cycle timing. A prime candidate for this coordination is ERK-1/MPK-1 signaling, which is important for proper germline architecture, oocyte size, and progression into both pachytene and diplotene (Church et al., 1995; Lee et al., 2007a; Lee et al., 2007b; Robinson-Thiewes et al., 2021). Furthermore, this signaling pathway has been implicated in regulating oocyte size and apical constrictions of apoptotic germ cells (Kohlbrenner et al., 2024). Understanding how ERK/MAPK signaling is affected by increased germline compartment number, and how defects in this signaling pathway affect germline architecture in apoptotic defective animals, will be the focus of future inquiry.

Apoptosis has been shown to play a plethora of roles during development and tissue homeostasis in animals (Fuchs and Steller, 2011; Ghose et al., 2018; Voss and Strasser, 2020). It can act as a quality control mechanism removing cells bearing defects caused by environmental stress, infection, or mitotic errors (Elmore, 2007; Luo et al., 2010; Norbury and Hickson, 2001; Raiders et al., 2018). On the other hand, apoptosis plays vital roles in eliminating specific populations of otherwise healthy cells to support tissue morphogenesis and architecture. The role of apoptosis in removing germline compartments containing DNA damage caused by environmental stress or meiotic failures is well documented in *C. elegans*; the persistence of such defective oogenic compartments is likely reflected in the ∼20% of baseline embryonic lethality observed at all timepoints in *ced-3(lf)* worms (Andux and Ellis, 2008; Raiders et al., 2018). The fact that loss apoptosis also leads to significant changes in germline architecture associated with defects in fertility and embryonic viability suggests that the role of apoptosis in the germline extends beyond ensuring nuclear quality control to an important contribution to germline architecture. Our findings suggest that these processes collaborate to maximize the number of viable fertilized embryos. How germline architecture and, by extension, germline compartment volume, is regulated is less well understood, but it is generally assumed that a delicate force balance between contractility of the actomyosin cortex that lines the rachis, and cytoplasmic streaming within this common cytoplasm is required for maintaining proper germline architecture and regulate oocyte size (Nadarajan et al., 2009; Priti et al., 2018; Rehain-Bell et al., 2017; Wolke et al., 2007). As expected, the increase in total compartment number due to a lack of apoptosis in *ced-3(lf)* animals leads to defects in germline architecture and thus oocyte size. We also note the increased nuclear localization of ANI-1 in “stacked” or “clustered” oocytes in animals lacking apoptosis; this change in the localization of a cortical contractility protein may affect the force balance among germline compartments. These defects however, are likely not due to decreased cytoplasmic streaming since flow speeds have been reported to be largely normal in *ced-3(lf)* animals (Wolke et al., 2007). Instead, *ced-3(lf)* germline defects may be caused by a combination of delayed entry into pachytene, postponed oocyte enlargement and thus less time for this process before detaching from the common cytoplasm during cellularization, and/or spatial limitation of oocyte enlargement due to progressive overcrowding of the germline with compartments that ordinarily would be removed via apoptosis.

Another developmental window in which apoptosis has been implicated in germline function is in aging worms past the normal window of fertility. As hermaphrodites age and sperm are depleted, the germline progressively atrophies (Kocsisova et al., 2025; Kocsisova et al., 2019). This process is accelerated by apoptosis (de la Guardia et al., 2016). Feminized worms (that do not produce sperm) aged for various durations before being provided sperm via mating exhibited an age-dependent reduction in embryonic viability that is significantly enhanced in the absence of apoptosis (Andux and Ellis, 2008). Interestingly, in this context as in our work, increased embryonic lethality correlated with decreased oocyte size, suggesting that both during the time-window of normal fertility and in aged virgin females, similar mechanisms for regulating oocyte quality via maintaining germline architecture are at work. These similarities raise the possibility that another biological role for germline apoptosis is to manage germline architecture and oocyte production in response to sperm availability, executing the dismantling of the germline, redirecting metabolic resources in the absence of sperm, and promoting proper germline architecture and oocyte size when sperm are provided via mating. It will be interesting to investigate in detail how germline architecture evolves in these conditions.

## Acknowledgements

We are grateful to the *Caenorhabditis* Genetics Center and Dr. Kacy Gordon for providing nematode strains, Bob Goldstein for sharing RNAi reagents, and the lab of Dr. Rob Dowen for their assistance with DIC imaging. We also thank Dr. Kacy Gordon for the thoughtful comments on the manuscript. We thank members of the Maddox labs for helpful discussion, and Noel Saydee, for arriving on time.

## Materials and Methods

### *C. elegans* strains and culture

*C. elegans* strains were maintained at 20°C using standard procedures (Brenner, 1974). The strains used in this work are listed in Table 1. MDX129 was generated by crossing MDX40 and MT1522. MDX155 was obtained by crossing UM208 and KLG049. MDX187 was created by crossing MDX155 and MT1522. MDX63 was generated by crossing MDX29 and LP193. KLG049 was a generous gift from Kacy Gordon and generation of the allele was initially described in (Roy et al., 2018).

**Table 1.**
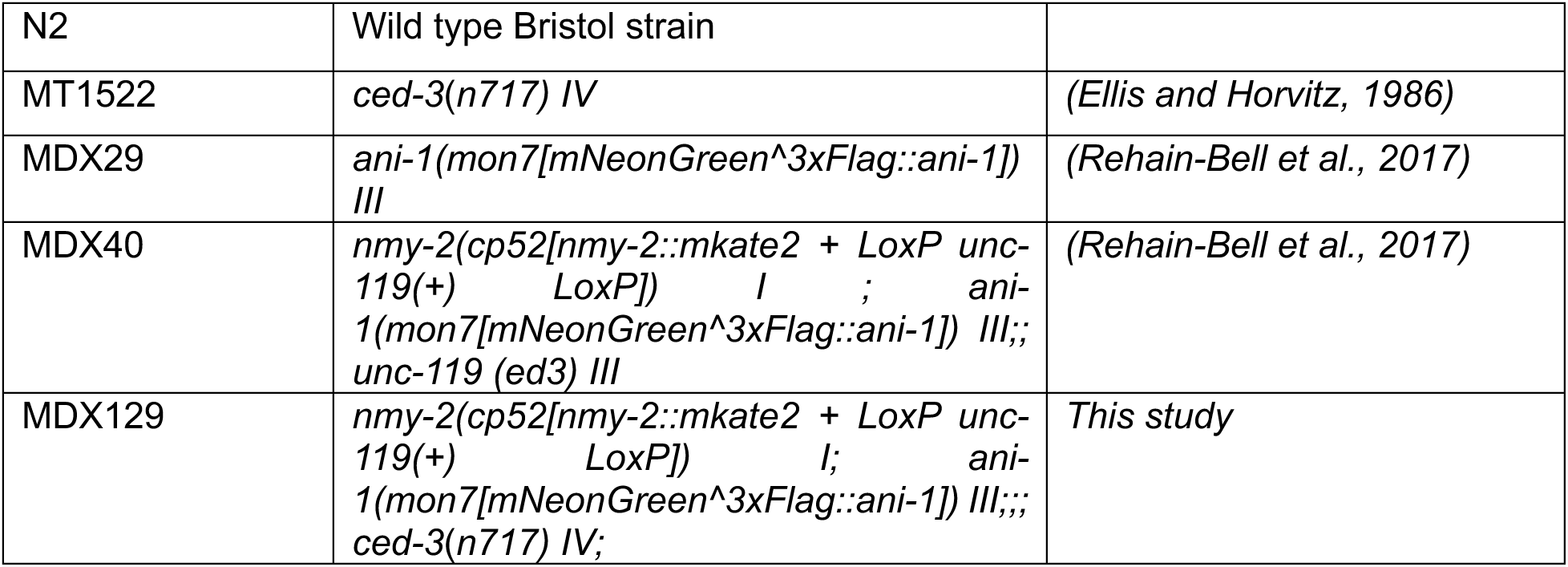
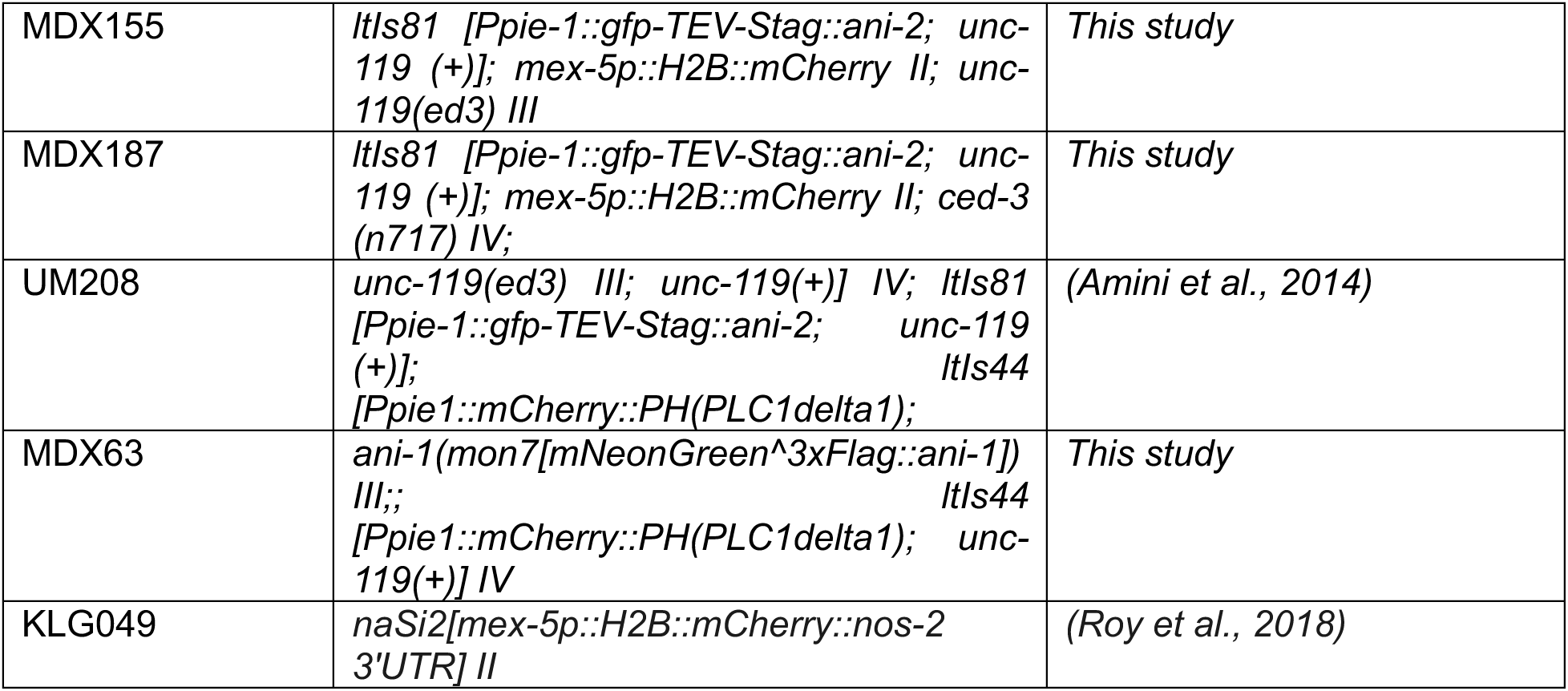

### RNA interference (RNAi)

RNAi was performed by feeding HT115 *E. coli* bacteria expressing double-strand RNA at 20°C for 24 hours as described previously (Kamath and Ahringer, 2003; Kamath et al., 2003). Individual bacterial clones were obtained from RNAi library (gift from Bob Goldstein, UNC) (Kamath and Ahringer, 2003).The targets of clones used in this study were confirmed by sequencing. RNAi depletions were monitored for different experiments at 24, 44, and 60 hours post L4.

### Synchronizing *C. elegans*

*C. elegans* synchronization was performed as previously described (Perry et al., 2023). L1 animals were monitored until L4 when they were collected and mounted for imaging of their germlines at 24, 44, and 72 hours post L4.

### Imaging *C. elegans* germlines and embryos

#### Imaging dish preparation

A mixture of 1% agarose and 5 mM levamisole in 50 ml of M9 buffer was heated in a conical flask until agarose was dissolved, briefly cooled, and poured, ∼3 ml quantities into 35 mm Mattek coverslip-bottom imaging dishes (*MatTek Corporation, Part No. P35G-0.170-14-C*). Dishes were cooled for 30 minutes in a hood, wrapped with parafilm, and stored at 4°C for use no longer than 4 months.

#### Live mounting for imaging

For all imaging experiments live hermaphrodites were picked at larval stage 4 (L4) and mounted into 35 mm Mattek agarose imaging dishes: With the 1% agar pad in a prepared imaging dish gently lifted, 0.8 µL of 5 mM levamisole in M9 buffer was pipetted onto the coverslip. For still imaging of live animals, 5 to 7 live hermaphrodites were gently placed onto the 0.8 µL spot using a platinum pick, and, after immobilization, positioned with an eyelash tool. The agar pad was gently repositioned in contact with the plate bottom to hold animals in place for live imaging via gentle compression.

DIC microscopy was performed on a Nikon SMZ-18 Stereo microscope equipped with a DS-Qi2 monochrome camera, at 67 ms exposure. The fluorescence imaging of live worms was performed using a 60X, 1.27 NA Nikon Water Immersion Objective on a Nikon A1R microscope with a Gallium arsenide phosphide photo-multiplier tube (GaAsP PMT) detector using NIS-elements. Approximately 35 optical sections, separated by 1 µm, were collected for Figure 4. All fluorescence imaging was performed by this method unless specified. Still images of germlines in Figure 3C and D were collected with a 10X, NA 0.30 objective on a Nikon A1R, 5 µm z-steps, and ∼7 total steps collected (only middle-section shown).

#### Imaging Embryos

L4 animals of indicated genotypes were grown for 24 or 44 hours on NGM plates. Animals were picked into a 2-5 µL droplet of M9 on the coverslip of a premade Mattek imaging dish with a 1% agarose pad (lift off as described above). A hypodermic needle was used to cut gravid adults near the uterus to release embryos into M9 buffer (0.85 M Na_2_HPO_4_, 0.4 M KH_2_PO_4_, 0.3 M NaCl, 1 mM MgSO_4_)). An eyelash tool was used to position embryos horizontally on Mattek imaging dishes. The 1% agar pad was slowly and gently re-positioned over the embryos and dishes were mounted on the stage for imaging. Images were acquired with a Hamamatsu Orca FusionBT camera on a Nikon Eclipse Ti2-FP, with a 20X, 0.75 NA Objective, with transmitted light, and the FITC channel via LED illumination.

For timelapse imaging, more M9 buffer was provided on the cover slip, plates were covered with designated lids, wrapped with thin pieces of parafilm, and acquisitions were collected at 4-hour intervals for 24 hours.

### Brood size analysis

Ten to twelve N2 and MT1522 hermaphrodites at larval stage 4 (L4) were placed onto NGM plates seeded with OP50 *E. coli*. The number of embryos laid by each animal was counted in the time windows of 0-24h, 24-48h and 48-60h, consecutively. Animals were moved to new NGM to fresh plates at each time point. The total number of hatched and unhatched embryos per adult at each timepoint were combined to determine the total 60-hour brood count.

For measuring embryonic viability, we allowed embryos to hatch and counted the number of live progeny for each adult ∼40 hours after assessing brood size. Embryonic viability was calculated at each timepoint as follows:

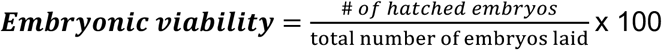

### Image Analysis

Image analysis was performed using the FIJI/Image J software (Schindelin et al., 2012). Unless specified otherwise, maximum intensity projections of z-sections were generated to show cortices, intercellular bridges, rings, rachis, and oogonia compartments starting at the distal tip through to the proximal germline arm.

Assessment of cell-cycle zones: The cell-cycle zones of oogonia were discerned based on chromatin morphology. Zones were defined as described (Lints and Hall, 2009): Mitotic (distal tip nuclei devoid of crescent shaped histones), Transition (chromatin occupying a crescent within the nucleus), Pachytene (“bowl-of-spaghetti” chromatin appearance), Diplotene/diakinesis (chromosome bivalents).

Rachis width was determined manually using Fiji (Schindelin et al., 2012) by measuring rachis diameter in various parts of the germline using ANI-2::GFP (MDX155, and MDX187) to mark the outline of the rachis lining. These measurements were used to calculate cross-sectional rachis area by approximating the rachis cross-sectional area to a circle using a custom R-script.

Germline compartment number was determined using automated segmentation of germline nuclei in the distal part of the germline before the bend in strains expressing H2B::mCherry (MDX155 and MDX187) using Cell-ACDC (Pachitariu and Stringer, 2022; Padovani et al., 2022). Segmentation results were curated manually to remove misannotated areas and include nuclei missed by the segmentation. For parts of the germline proximal to the bend the segmentation failed, and nuclei were identified manually using Fiji.

To measure germline compartment cross-sectional area, compartments were approximated to be rectangles in the strain MDX63 expressing mCherry::PH(PLC1delta1) to label the plasma membrane. Animals were treated with control or *ced-3(RNAi)* for 44 hours and length and width of the germline compartments were measured by placing a point on apical and basal as well as each lateral membrane of a compartment respectively, using the multipoint tool in FIJI. Cross-sectional area was estimated by calculating the distance between the 2 pairs of points and using these values as width and length measurements to calculate the area of the rectangle using a custom Python script.

For all measurements, the basal tip of the bend of the germline was used as a reference point to position the individual measurements along the length of the germline, by calculating the distance between the reference point and the position of the center of the measured objects.

Measurements distal to the bend were assigned a negative positional value while proximal measurements have positive positional values, to plot the measurements as developmental time courses in the same graph. For graphical depiction of developmental time courses of both rachis area and germline compartment number, measurements were binned into 10 µm bins based on their position relative to the reference point in the germline bend using custom scripts in R and Python. Mean values and 95% confidence intervals were calculated and plotted for each bin using Microsoft Excel.

Embryo length along the A-P axis was measured using Fiji after extruding embryos from the animal.

### Figures

Figures were assembled in Microsoft PowerPoint and Adobe Illustrator. Prism (GraphPad), Microsoft Excel, and Seaborn Python data visualization library (Waskom, 2021) were used for data visualization.

### Statistical analysis

Statistical testing and comparisons were performed using Prism (GraphPad) or Microsoft Excel as described in corresponding figure legends. A p-value of less than 0.05 was considered significant. Statistical significance and respective p-values are shown in all figures and corresponding figure legends. All error bars represent standard deviations. Sample size (n) is indicated in the text or the figure legend.

---

**Supplemental Figure 1.**
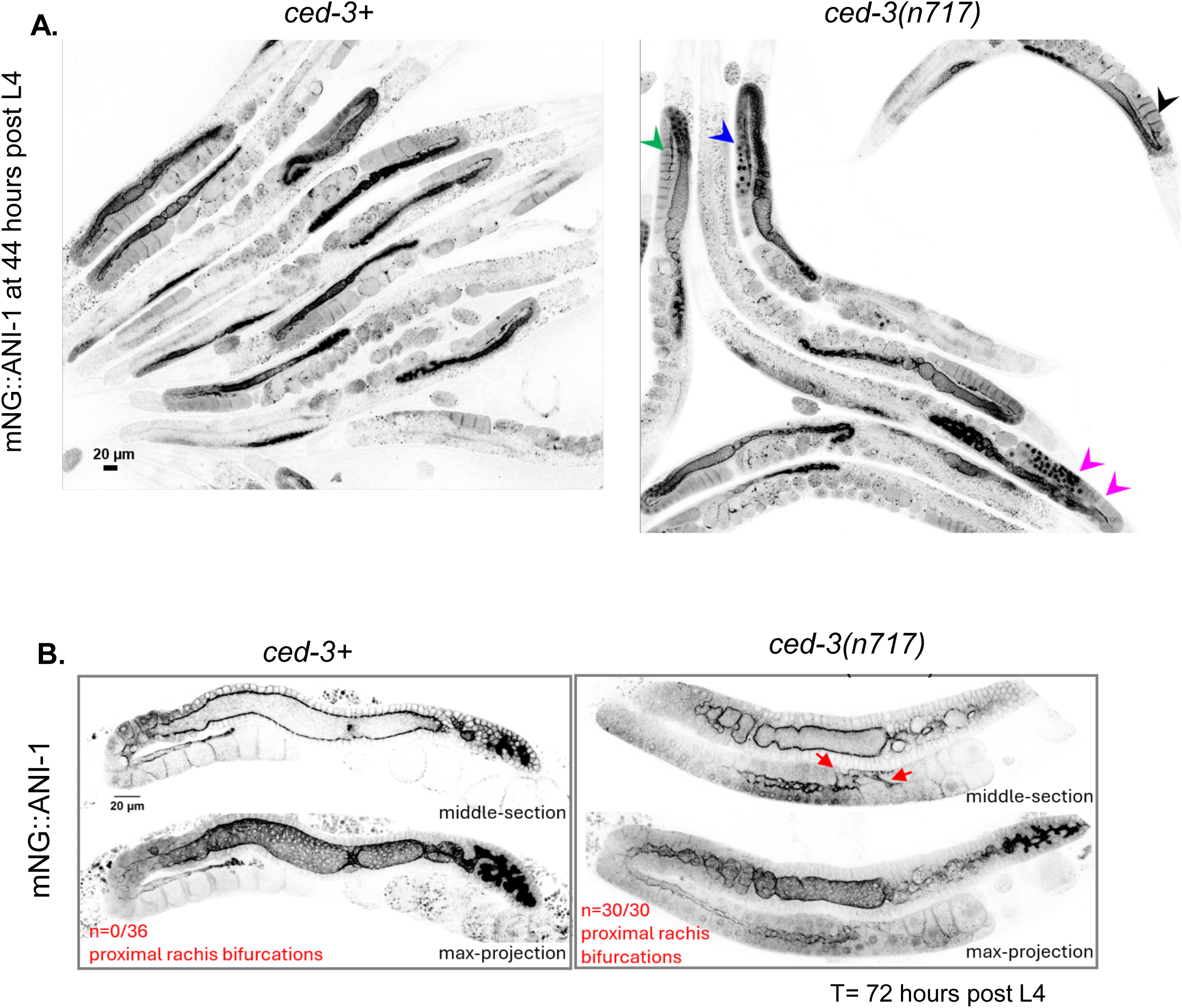
Brood size and germline architecture defects intensify over the reproductive life of *ced-3*(*lf*) hermaphrodites. **A.** Representative germline morphologies observed in *ced-3*(*lf*) animals at 44h post L4 are shown. Arrowheads point to normal (Black), clustered (Blue), stacked (Green), clustered germ compartment nuclei and stacked oocytes (Magenta). **B.** Representative images showing germline morphology via mNG::ANI-1, at post-fertility age (L4+72h) in the indicated genotypes, red arrows point to bifurcations of the rachis observed in *ced-3* mutant germlines.

